## Supplementary Information for "Detection and skeletonization of single neurons and tracer injections using topological methods"

### Supplementary Methods

**Implementation in the discrete setting** The following provides a more detailed description of how the persistence-guide Morse-based graph framework is implemented. For even more specifics, please refer to [20] for an introduction into Discrete Morse Theory and to [19] for more details on the algorithm we implement.

**Algorithm 1** $G$ = `DiMorSC` (K, $\rho$, $\tau$)

1: Persistence Computation
- Compute persistence pairings induced by lower-star filtration of K with respect to -$\rho$

2: Obtain Simplified Discrete Gradient Vector Field
- Initialize trivial vector field
- For each persistence pair, perform cancellation if possible and persistence $\leq \tau$

3: Collect Output
- compute the 1-unstable manifold of each critical edge with persistence $> \tau$

4: **return** union of 1-unstable manifolds

The Discrete Morse algorithm is traditionally given a triangulation K of the domain and a density function $\rho$ given at the vertices of K. Instead of a triangulation, we take K to be the 2-skeleton(vertices, edges, and squares) of the cubical complex of the domain. This does not change any part of the algorithm and reduces computation time. Additionally, the user provides the algorithm with a persistence threshold $\tau$.

**Step 1** The first step of the algorithm is to compute the persistence pairings P(K) by the lower star filtration of K with respect to -$\rho$. In our implementation, we use DIPHA[63] to compute persistence because it is a distributed persistent homology algorithm that we found minimizes computation time.

**Step 2** The second step of the algorithm is to compute the discrete gradient vector field. As shown in [19], all that is needed is to calculate the spanning forest that is made up of all negative edges (edges that are paired with a vertex in P(K)) with persistence less than or equal to $\tau$. Positive edges (edges paired with a square) and edges with persistence greater than $\tau$ are not part of the spanning forest. No explicit discrete gradient vector field needs to be computed nor maintained. This step takes linear time once the persistence pairings are computed in Step 1.

**Step 3** The third step of the algorithm is to compute the 1-unstable manifold of each critical edge. As shown in[19], for each edge, the 1-unstable manifold is equivalent to the union of the edge with the paths from both vertices to the sink of their corresponding tree in the spanning forest computed in Step 2. The union of all 1-unstable manifolds is outputted by the algorithm. Please note that the 1-unstable manifold of -$\rho$ is equivalent to the 1-stable manifold of $\rho$.

**Initial vector-based Morse graph simplification.** Given the Morse skeleton graph computed in Step 2 of the DM-Skeleton pipeline, at the beginning of Step 3 of our pipeline, we can perform a flow-vector based graph simplification to remove branches misaligned with underlying flow vectors. In particular, we first estimate flow-vectors to get a sense of which direction true neuron branches will flow. To do this, we use a weighted principal component analysis [60]. In standard principal component analysis, the principal component represents the direction which explains the most variance of a collection of data points. With weighted principal component analysis, the principal component represents the direction which explains the most variance of the weights of data points. For each vertex in the Morse graph output, we compute this flow-vector for a cubic neighborhood, assigning each point in the neighborhood a weight equal to its intensity. We then apply Gaussian diffusion to all computed flow-vectors for the sake of smoothing. Note that the flow-vector estimation is only carried out for nodes in the Morse skeleton graph.

Next, each path from non-degree 2 to non-degree 2 node in the Morse skeleton graph is computed. Then, for each vertex $v$ in each path $s$, we estimate a path-vector at $v$ w.r.t. $s$ by taking the difference between the two vertices that are 4-hops from $v$ in the path. We compute the cosine between this path-vector and the estimated flow-vector at $v$ and consider this to be the vector score at $v$. The closer the value is to 1, the more the flow-vectors and path-vectors are aligned, meaning the Morse graph skeleton is more aligned with the estimated flows. On the other hand, the closer the value is to 0, the less the vectors are aligned, indicating the Morse graph skeleton is perhaps significantly deviating from the estimated flows. Note that for each vertex $v$ with degree greater than two, $v$ receives a score for each path it is a part of.

In addition to these vector scores, a capped intensity value is calculated for each vertex in the Morse skeleton graph. This is simply the minimum of the corresponding voxel value of the vertex and a user-provided value. For our experiments, the value provided is 1.

Finally, a score is computed for each path using the vector scores and capped intensity values. Specifically, for each vertex $x$ in each path $s$, let $c(x)$ return the capped intensity value of $x$, and let $v(x)$ return the vector score of $x$ w.r.t. $s$. The score of the path is equal to $\int_s c(x)(\alpha + v(x))ds$ divided by the length of $s$. $\alpha$ is a user provided weight parameter (a value of zero is used in our experiments).

Once scores are computed for each path, the simplification process begins. Only paths below a user-provided threshold are removed. In increasing score order, paths are removed if doing so does not change the number of connected components in the Morse graph skeleton. If a path is removed, it is possible that paths with lower scores that were not removed will now not alter connectivity if removed, and thus are again tested for removal. The simplified Morse skeleton graph is then fed into a weighted shortest spanning forest algorithm.

**Weighted shortest spanning forest algorithm** Given a graph $G = (V, E)$ where each node $v \in V$ is a point in $\mathbb{R}^3$, let $R \subset V$ with $R = \{v_0, ..., v_m\}$ denote the set of roots for spanning forests. We then compute a weighted shortest-spanning forest for $G$ as follows: For any edge $(u, v) \in E$, its weight is defined as $w(u,v) = \frac{2d(u,v)}{\rho(u)+\rho(v)}$, where $d(u,v)$ is the Euclidean distance between $u$ and $v$, and $\rho : V \to \mathbb{R}$ is the density map. Using these weights (as distances), the algorithm computes the shortest path distances between each root in the root-set $R$ and nodes in $V$. Then for each root $r \in R$, its shortest path tree $T_r$ is spanned by all nodes whose shortest path distance to $r$ is smaller than that to any other node in $R$ (ties are broken arbitrarily).

**Score initialization and smoothing** We now describe how to compute a score $\overline{s}(v)$ for all tree nodes in a given tree $T$ (from the spanning forest), so as to carry out the tree simplification procedure as described in Methods section.

Consider a specific tree $T_r$ from the weighted shortest-path forest, with root $r \in R$. We first calculate two temporary scores carrying different types of information for each tree node $v$ in $T_r$: one is a density score $dscore(v)$ based on density information, and the other one is a vector score $vscore(v)$ based on directional information. The weighted score $s$ of each tree node $v$ is the weighted sum of these two temporary scores, i.e., $s(v) = \alpha \cdot dscore(v) + (1-\alpha) \cdot vscore(v)$ with $0 \leq \alpha \leq 1$. Empirically, a large $\alpha$ is used because density is the major indicator for false positives. The details for computing $dscore$ and $vscore$ are stated in the following sections. After the weighted score $s$ is computed for all tree nodes, we further smooth this score to obtain the final score $\overline{s}$.

**(1). Calculate density scores for tree/graph nodes.** The density score $dscore(v)$ is calculated based on voxel density. For each voxel, we first find its nearest neighbor in the set of tree nodes $V$ and associate it with the nearest neighbor. To avoid associating a voxel to a tree node far away, we set an upper bound $\beta_s$ , thus a voxel will not be associated with any tree node if its distance to the nearest neighbor exceeds this upper bound. Empirically, on both injection tracer and fMOST single-neuron datasets, the default upper bound $\beta_s$ is set as the distance between adjacent brain slices (i.e., 1μm for the fMOST dataset and 50μm for the STP dataset). If a voxel has multiple nearest neighbors, we break the tie arbitrarily. The density score $dscore$ of a tree node $v \in V_s$ is simply the sum of density values of all associated voxels, i.e.,

$$dscore(v) = \sum_{x \in I, d(x,v)=d(x,V_s), d(x,v) \leq \beta_s} \rho(x),$$

where $I$ denotes the set of all voxels, $\rho$ is the density map and $V_s$ is the node set of the simplified tree (Fig. 3b).

**(2). Calculate vector scores for tree nodes.** Vector scores have already been computed as described in Initial vector-based Morse graph simplification earlier.

**(3). Score smoothing to obtain final score $\overline{s}$.** Recall that we obtain a combined weighted score $s(\cdot)$ from density scores and vector scores. This weighted score will be further smoothed before performing tree simplification to remove local noise. In particular, the score of a tree node $v$ will be smoothed by averaging those of its neighbors within $k$-hops (i.e., connected via at most $k$ tree edges to $v$). We further restrict only to neighbors that are ancestors or descendants of $v$ given the root $v_0$: intuitively, we wish to consider only neighbors of $v$ along the "same" neural branch. An example with $k = 3$ is shown in Fig. 3c. The final score of tree node $v$ is $\overline{s}(v) = \frac{1}{|H_v^k|} \sum_{x \in H_v^k} s(x)$, where $H_v^k$ denotes the set of $v$'s ancestors and descendants (including $v$ itself) within $k$ hops. In practice, maximum hop $k$ is set to 10 for both tracer injection and fMOST datasets.

#### Evaluation metrics

**Precision, Recall and F1-score for evaluating fMOST results.** For fMOST data, the single neuron reconstruction data were provided as groundtruth. The precision and recall metrics calculated for evaluating fMOST results depend on True Positives (TP), False Positives (FP) and False Negatives (FN). All the outputs were discretized before computing these metrics. We broke any segment with a length greater than 2 pixels so that segment lengths are all roughly 1 pixel. Then, for each node $v$ from the discretized skeletonization, we labelled $v$ as either TP or FP. $v$ is considered as TP if its nearest neighbor in the human annotation is within $4\mu m$. Otherwise, $v$ is a FN. Similarly, a node in the annotation is a FN if there is no predicted node within $4\mu m$. Precision and recall are then routinely computed as $Precision = \frac{TP}{TP+FP}$ and $Recall = \frac{TP}{TP+FN}$. The F1-score is the harmonic mean of precision and recall, i.e., $F_1 = \frac{2 Precision \cdot Recall}{Precision + Recall}$. The parameter $4\mu m$ was predetermined based on similar reasons as in [50]. The thickness of dendrites near the soma is around $4\mu m$ and the curve of F1-score against this parameter (Supplementary Fig. 1) also supports the choice. The F1-score is stabilized when the parameter reaches 4μm.

**Region analysis** The brain-wide inter-regional connectivity information was obtained through region analysis on the whole brain tracing data. The 3D volume of the data was registered with the Allen Mouse Brain Atlas at 10 μm resolution [61]. Based on the atlas, the 3D density fields (proofread STP dataset and projected summarizations) were segmented into individual brain regions. Regions were considered "covered" when the total weights of the voxels in these regions were greater than zero. The covered brain regions were ranked based on the total weights of the voxels inside. The two hemispheres were separated and the above described region analysis was performed on each side in addition to the whole-brain analysis. To quantitatively evaluate performance, we calculate a coverage rate for each method on the two hemispheres, and the entire brain. For a given method, the coverage rate is equal to the sum of the weights for regions in which the method's output covers divided by the sum of weights for all regions.

| | Method | Precision | Recall | F1 | Proofreading time | Total process length (cm) | Normalized proofreading (sec / cm) |
|---|---|---|---|---|---|---|---|
| Neuron1 | GTree | 0.846 | 0.906 | 0.875 | - | 0.500 | - |
| | GTree-a1 | 0.953 | 0.940 | **0.946** | 23m40s | | 2839.06 |
| | GTree-a2 | 0.954 | 0.929 | 0.941 | 21m40s | | 2599.14 |
| | DM | 0.900 | 0.940 | 0.920 | - | | - |
| | DM-a1 | **0.957** | 0.923 | 0.940 | **15m43s** | | **1885.37** |
| | DM-a2 | 0.948 | **0.941** | 0.944 | 15m56s | | 1911.37 |
| Neuron2 | GTree | 0.749 | 0.750 | 0.750 | - | 0.508 | - |
| | GTree-a1 | 0.818 | 0.748 | 0.781 | 20m00s | | 2399.20 |
| | GTree-a2 | 0.807 | 0.756 | 0.780 | 21m40s | | 2599.14 |
| | DM | 0.865 | 0.749 | 0.803 | - | | - |
| | DM-a1 | 0.866 | **0.773** | 0.817 | **16m00s** | | **1919.36** |
| | DM-a2 | **0.944** | 0.771 | **0.849** | 16m21s | | 1961.35 |
| Neuron3 | GTree | 0.621 | 0.729 | 0.670 | - | 0.640 | - |
| | GTree-a1 | 0.781 | 0.744 | 0.762 | 23m21s | | 2801.07 |
| | GTree-a2 | 0.726 | 0.746 | 0.736 | 22m45s | | 2729.09 |
| | DM | 0.890 | **0.789** | 0.836 | - | | - |
| | DM-a1 | **0.939** | 0.771 | 0.847 | 18m20s | | 2199.27 |
| | DM-a2 | 0.925 | **0.789** | **0.852** | **17m46s** | | **2131.29** |

**Supplementary Table 1** | Proofreading is done on GTree and DM-Skeleton (abbreviated as DM in this table) outputs on all three fMOST neuron regions. We further computed the evaluation metrics on those proofread results with the original ground truth. The suffixes "a1" and "a2" represents the two annotators for proofreading. The F1-scores are improved after proofreading. The proofreading time spent on the DM-Skeleton method is also significantly shorter.

| Simplification \ Persistence | 128 | 256 | 512 | 768 | 1024 |
|---|---|---|---|---|---|
| 0 | 0.352 | 0.368 | 0.468 | 0.503 | 0.445 |
| 0.05 | 0.899 | 0.911 | 0.912 | 0.828 | 0.639 |
| 0.10 | 0.909 | 0.918 | 0.910 | 0.825 | 0.627 |
| 0.15 | 0.910 | 0.918 | 0.907 | 0.818 | 0.590 |
| 0.20 | 0.912 | **0.920** | 0.899 | 0.791 | 0.584 |
| 0.25 | 0.902 | 0.910 | 0.889 | 0.788 | 0.583 |
| 0.30 | 0.897 | 0.902 | 0.872 | 0.776 | 0.582 |

**Supplementary Table 2** | This table shows the F1-score of DM-Skeleton outputs with different simplification and persistence thresholds. The F1-score is calculated based on the region of neuron1. The result of the optimal F1-score uses persistence threshold = 256 and simplification threshold = 0.2. The same thresholds (256, 0.2) are applied to the other two regions.

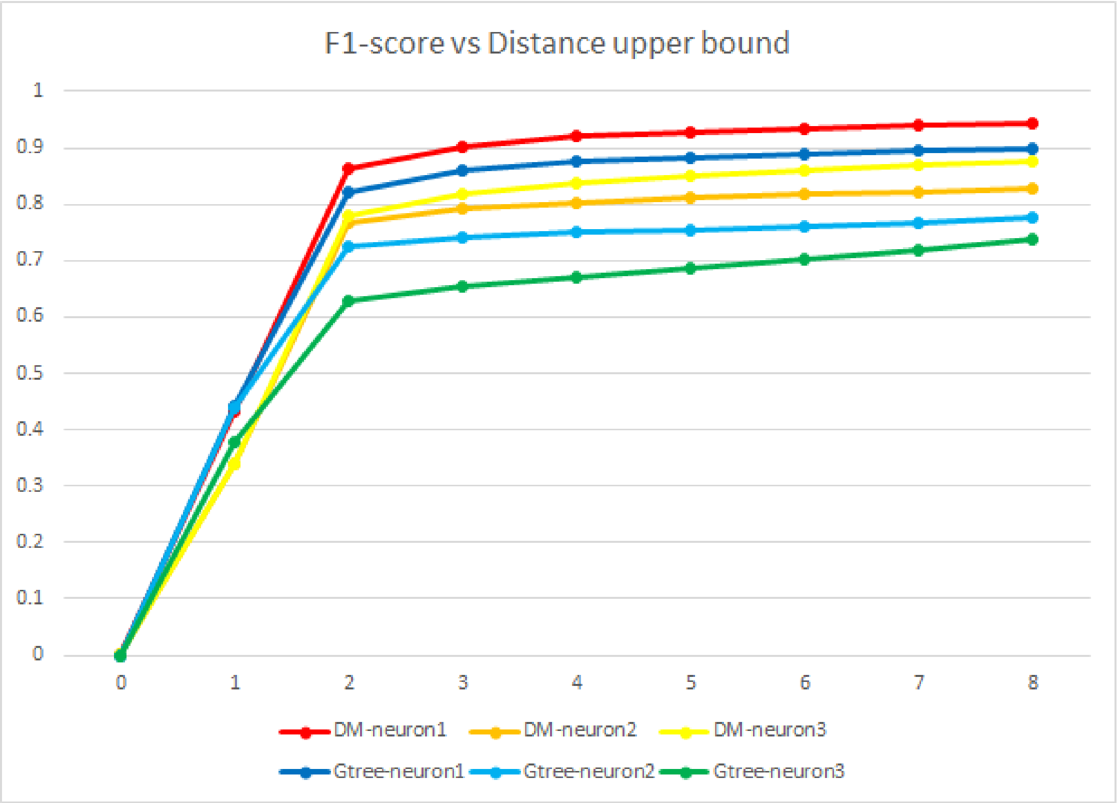


**Supplementary Fig. 1** | F1-scores vs different distance bounds. The F1-scores for both DM-Skeleton and GTree outputs start to stabilize when the upper bound becomes larger than 4 µm. Therefore 4µm is a reasonable choice for the bound.

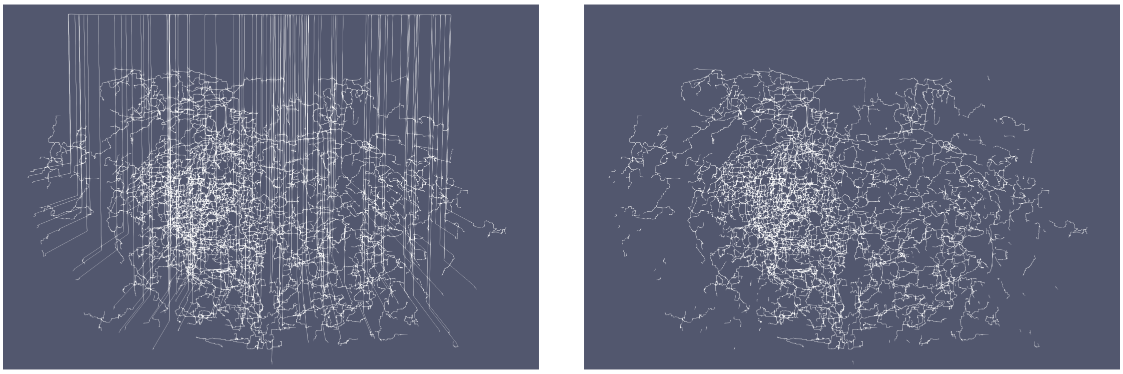

**Supplementary Fig. 2** | On the STP dataset, due to the degenerated gradient on the background pixels, there are redundant edges going to the boundary (on the left). Those edges can be filtered out by a small density threshold (result on the right) since the pixel value of the background is zero after pre-processing.

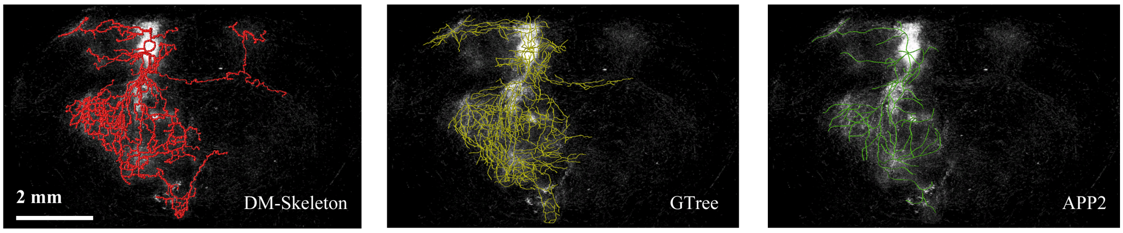


**Supplementary Fig. 3** | Summarization results of DM-Skeleton method (red), GTree (yellow) and APP2 (green).

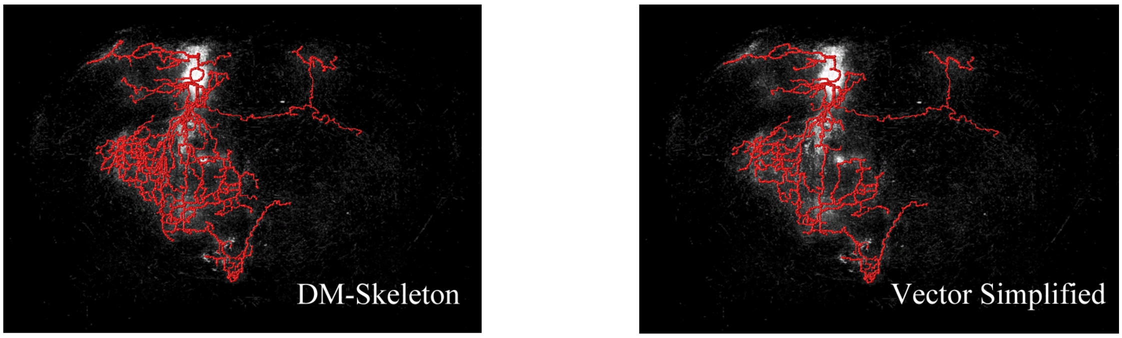


| Method | Top 5 non-injection regions in proofread brain (rank) | | | | | Other selected regions | | Contralateral coverage | Ipsilateral coverage | Overall coverage |
|---|---|---|---|---|---|---|---|---|---|---|
| | CP(i) | MOs(i) | SSp(i) | SC(i) | APN(i) | MOp(c) | MOs(c) | | | |
| Proofread | 13.44% (1) | 9.56% (2) | 4.83% (3) | 3.76% (4) | 3.05% (5) | 1.34% | 0.63% | - | - | - |
| DM | 15.08% (1) | 8.08% (2) | 4.24% (3) | 3.75% (5) | 3.98% (4) | 1.48% | 0.26% | **57.99%** | **96.24%** | **93.34%** |
| DM (vector) | 15.37% (1) | 6.47% (2) | 3.07% (6) | 3.80% (5) | 3.95% (3) | 1.45% | 0.33% | 48.34% | 93.96% | 90.50% |

**Supplementary Fig. 4** | Summarization results of DM-Skeleton (abbreviated as DM in the table for better presentation) method, and the version after vector simplification. The coverage percentages are shown in the table. We can see that the vector simplification can provide a clearer structure without much loss of coverage. All the selected regions are still covered by the DM-Skeleton output after vector simplification, but the ranking of regions is slightly distorted.

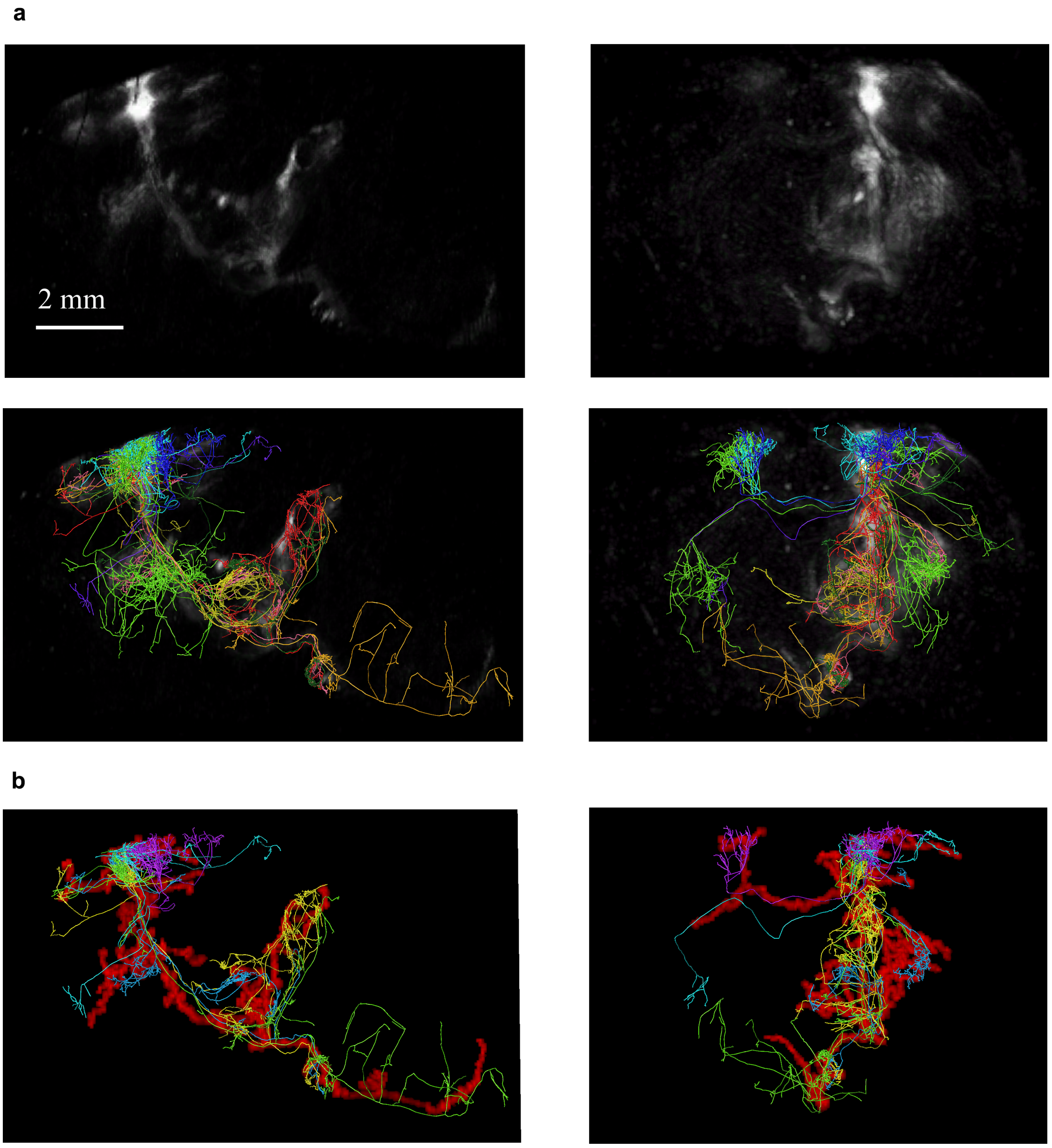


**Supplementary Fig. 5** | Mouselight superposition. **a,** The selected 11 mouselight neurons are superposed with the proofread volume and compared in 2 angles. We mapped the volume to atlas space and slightly raised the brightness and contrast for easier comparison. **b,** The neurons are also superposed with the DM-Skeleton output. To see the clear contour of DM-Skeleton summarization, we only selected a representative subset of those 11 neurons which can show the shape with lower complexity.
